# Atrazine and amphibians: Data re-analysis and a summary of the controversy

**DOI:** 10.1101/164673

**Authors:** Jason R. Rohr

## Abstract

The herbicide atrazine is one of the most commonly used, well studied, and controversial pesticides on the planet. Much of the controversy involves the effects of atrazine on wildlife, particularly amphibians and their non-infectious and infectious diseases, including diseases caused by trematode infections. Here I re-analyze data from authors that were funded by Syngenta Crop Protection, Inc., the company that produces atrazine, and show that even these authors revealed that increasing concentrations of atrazine applied to outdoor mesocosms increases the population growth rate of snails that can transmit trematode parasites to amphibians. These researchers missed this finding in their data because they never calculated population growth rates for the snail populations before they reached a carrying capacity or crashed. These results demonstrate that both Syngenta-funded and non-Syngenta-funded researchers have provided evidence that ecologically relevant concentrations of atrazine are capable of increasing snail populations. Given the controversy surrounding the effects of atrazine on amphibians, I follow this re-analysis with a timeline of some of the most salient events in the history of the atrazine-amphibian controversy.

## Introduction

The herbicide atrazine [2-chloro-4-(ethylamino)-6-(isopropylamino)-S-triazine] is one of the most widely studied, commonly used, and controversial pesticides on the planet. A search for the term “atrazine” in the search engine Web of Science (conducted on 11/17/2016) produced 11,203 studies. Atrazine was the most commonly used pesticide in the US before it was recently surpassed by the herbicide glyphosate (Roundup®), which happened because of the advent of genetically modified crops (Grube et al. 2011). Syngenta Crop Protection, Inc., the company that produces atrazine, earns approximately $2.5 billion annually from its selective herbicides, of which atrazine is their leading product. Because of it heavy use, as well as its persistence and mobility, atrazine is one of the most common chemical contaminants of freshwater and thus is regularly found in habitats where many freshwater vertebrates, such as fish and amphibians, develop (Rohr et al. 2003, Knutson et al. 2004). Consequently, there has been considerable interest in the effects of atrazine on freshwater vertebrates (e.g. Solomon et al. 2008, Rohr and McCoy 2010b), particularly amphibians because of their permeable skin and global declines (Rohr et al. 2008a, Wake and Vredenburg 2008, Rohr and Raffel 2010, Liu et al. 2013, Raffel et al. 2013, Rohr et al. 2015).

Research on the effects of atrazine on amphibians, however, has been highly contentious. Much of the controversy on atrazine and amphibians involves the effects of atrazine on their non-infectious diseases, such as disruption of the function and development of their endocrine and reproductive systems, and infectious diseases, such as trematode infections. Authors funded by Syngenta Crop Protection have argued that atrazine does not increase populations of snails that can transmit trematodes to amphibians (Baxter et al. 2011), whereas non-Syngenta-funded authors provide data suggesting that atrazine can increase snail populations (Rohr et al. 2008). Here, I briefly review the effects of atrazine on amphibians. I then re-analyze the snail data provided by authors that were funded by Syngenta Crop Protection, Inc. (Baxter et al. 2011). Given the controversy surrounding the effects of atrazine on amphibians, I then follow this reanalysis with a timeline of some of the most salient events in the history of the atrazine-amphibian controversy.

## Background on the Effects of Atrazine on Amphibians

Atrazine has a variety of effects on freshwater organisms, including fish and amphibians. For example, atrazine has been reported to affect amphibian behaviors crucial for foraging, predator avoidance (Rohr et al. 2003, 2004, Rohr et al. 2009), and desiccation resistance (Rohr and Palmer 2005, 2013). It also impacts growth and timing of metamorphosis (Larson et al.1998, Allran and Karasov 2000, 2001, Boone and James 2003, Rohr et al. 2004, Forson and Storfer 2006a, Forson and Storfer 2006b).

Given considerable interests in the role of physiology to vertebrate survival and conservation (Martin et al. 2010, Rohr et al. 2013b, Madliger et al. 2016), there have been numerous studies on the effects of atrazine on physiology. For example, several studies have investigated atrazine as an ‘infodisruptor’, defined as a chemical contaminant that disrupts communication within or among organisms, including contaminants that breakdown or interfere with detection or production of chemical signals between senders and receivers or those that affect cell-to-cell communication within organisms (e.g. endocrine disruptors)(Lurling and Scheffer 2007, Rohr et al. 2009). Several studies have shown that atrazine can reduce chemical detection of cues from predators and mates (Moore and Waring 1998, Tierney et al. 2007, Ehrsam et al. 2016) and others have shown that it can affect cell-to-cell communication by altering hormones, such as stress hormones (Gabor et al. 2016, McMahon et al. 2017), thyroid hormones (Larson et al. 1998), and sex hormones (Hayes et al. 2003, Hayes 2003). Given that much of the controversy regarding atrazine and amphibians involves its effects on amphibian sex hormones and gonadal development, this topic will be discussed in more detail in the “A Timeline of the Atrazine-Amphibian Controversy” section.

Interest in chemical contaminants causing non-monotonic dose-responses (those with a change in the direction of the slope) (Welshons et al. 2003, McMahon et al. 2011, Vandenberg et al. 2012, McMahon et al. 2013) has triggered several researchers to examine whether atrazine causes linear or non-linear dose responses. Researchers have detected non-monotonic dose responses on several amphibian hormones, including corticosterone, thyroid hormone, and sex hormones (Larson et al. 1998, Hayes et al. 2003, Hayes 2003, Fan et al. 2007, McMahon et al. 2017). Atrazine also regularly has non-monotonic effects on the timing of metamorphosis (Rohr and McCoy 2010b). Although responses on other endpoints have not produced non-monotonic dose responses as regularly, they have regularly produced logarithmic dose responses, where the greatest change in response occurs at low exposure concentrations (Rohr et al. 2004, Rohr et al. 2006b, McMahon et al. 2013, Rohr et al. 2013c), supporting the potency of low concentrations of atrazine.

Given that many factors have been documented to affect amphibian diseases that have been linked to amphibian declines (Li et al. 2013, Liu et al. 2013, Rohr et al. 2013b, McMahon et al. 2014, Venesky et al. 2014b), interest has grown in the role that atrazine might have on amphibian immunity and infections. Much of this interest accelerated in 2002 after Kiesecker (2002) revealed that atrazine exposure was associated with reduced amphibian immunity and increased trematode infections and the limb malformations they cause. Since then, Rohr and colleagues have found additional support for the immunosuppressive effects of atrazine and increases in trematode infections (Rohr et al. 2008b, Rohr et al. 2008c, Raffel et al. 2009, Schotthoefer et al. 2011, Rohr et al. 2015). Additionally, they showed that atrazine increases exposure to trematodes by reducing phytoplankton, which reduces shading and increases the abundance of periphyton, the food source for snails, which are the intermediate host of trematodes (Rohr et al. 2008c, Raffel et al. 2010, Staley et al. 2010, Staley et al. 2011, Halstead et al. 2014). Atrazine exposure, either alone or in mixtures with other chemicals, has also been associated with reduced immunity and increased amphibian viral, flatworm, and roundworm infections (Gendron et al. 1997, Forson and Storfer 2006a, Forson and Storfer 2006b, Hayes et al. 2006, Kerby and Storfer 2009, Koprivnikar 2010). Recently, atrazine exposure was shown to reduce tolerance of chytrid fungal infections that are associated with worldwide amphibian declines (Rohr et al. 2013c). Tolerance is defined as the ability of hosts to reduce damage caused by parasites (Rohr et al. 2010, Sears et al. 2013, Sears et al. 2015). In a 2010 review (Rohr and McCoy 2010b), atrazine exposure was associated with a reduction in 33 of 43 immune function endpoints and with an increase in 13 of 16 infection endpoints. These numbers were an underestimate (Langerveld et al. 2009) and have increased since this review was published (e.g. Koprivnikar 2010, Rohr et al. 2013c).

The documented positive relationship between biodiversity and ecosystem functions and services, such as pest and disease control, primary production, and clean water (Dobson et al. 2006, McMahon et al. 2012, Staley et al. 2014, Venesky et al. 2014a, Civitello et al. 2015, Cohen et al. 2016, De Laender et al. 2016) and the importance of indirect effects of chemicals mediated by species interactions (Rohr et al. 2006a, Clements and Rohr 2009, Halstead et al. 2014, Douglas et al. 2015, Staley et al. 2015) has prompted several researchers to study the effects of atrazine on freshwater communities containing amphibians rather than on isolated amphibian species (de Noyelles et al. 1989, Boone and James 2003, Rohr and Crumrine 2005, Rohr et al. 2008c, Halstead et al. 2014). Many of these studies report alterations of amphibian growth and abundance that seem to be caused by atrazine•induced changes in photosynthetic organisms. At ecologically relevant concentrations, atrazine is expected to have a bevy of indirect effects by altering the abundance of phytoplankton, macrophytes, (Herman et al. 1986) and photosynthetic and non-photosynthetic organisms in periphyton (Staley et al. 2010, Staley et al. 2011, Staley et al. 2012, Staley et al. 2015), the latter of which is a primary food source for many tadpole species. However, few of the studies focusing on atrazine and freshwater communities containing amphibians distinguish between direct and indirect effects of atrazine.

Although atrazine generally does not directly cause amphibian mortality at ecologically relevant concentrations (Solomon et al. 2008, Rohr and McCoy 2010b), there are some studies that suggest that it might increase mortality through indirect effects, such as those described in the infectious disease section above. Others suggest that there might be delayed or persistent effects of atrazine on behavior and physiology that can increase mortality risk (Storrs and Kiesecker 2004, Rohr and Palmer 2005, Rohr and McCoy 2010b, Rohr and Palmer 2013).

One of the biggest concerns regarding the effects of atrazine on amphibians is that atrazine regularly interacts with other stressors commonly experienced by amphibians, either additively or synergistically. For example, particular climatic conditions, such as increased drying (Rohr et al. 2004, Rohr and Palmer 2005, 2013) and particular biotic conditions, such as parasitism (Rohr and McCoy 2010b) and predation risk (Rohr and Crumrine 2005, Ehrsam et al. 2016) can be worsened by atrazine. Atrazine also additively or synergistically interacts with other common agrochemicals (Rohr et al. 2008c, Halstead et al. 2014). The exception is that global warming will accelerate amphibian development and thus reduce aquatic exposure to atrazine (Rohr et al. 2011). Clearly, there is a need to understand how climate change will affect exposure and toxicity of atrazine and other chemical contaminants (Rohr et al. 2013a, Landis et al. 2014). Nevertheless, interactions among chemical contaminants or between chemical contaminants and non-chemical stressors are unfortunately rarely considered in most ecological risk assessments of chemicals (Rohr et al. 2016, 2017).

## Methods

I extracted the data from Figure 5 of Baxter et al. (2011), which include authors who have historically been funded by Syngenta Crop Protection, Inc. In this experiment, various concentrations of atrazine were applied to outdoor mesocosms containing natural algae, zooplankton, and snails, and snail abundance was tracked through time. Given that initial levels of phosphorous and nitrogen could be depleted through time and were not replenished, these essential elements could become limited and cause algal and snail populations to reach carrying capacities or crash. Hence, I calculated the population growth rate of snails in each tank until an apparent carrying capacity or population crash occurred. Thus, growth rates were calculated for the exponential phase of population growth only. The 30 μg/L treatment was excluded because it did not show the same blocking patterns in dissolved oxygen as the other treatments (see Baxter et al. 2011 Table 3, Rohr et al. 2012). I then plotted the population growth rate estimates against log atrazine concentration.

## Results

Although not reported in the Baxter et al. (2011) paper, there was a clear increase in snail population exponential growth rates as atrazine concentration increased (Fig. 1). These researchers missed this finding in their data because they never calculated population growth rates for the snail populations before they reached a carrying capacity or crashed. These results demonstrate that both Syngenta-funded (Herman et al. 1986, Baxter et al. 2011) and non-Syngenta-funded researchers (Kiesecker 2002, Rohr et al. 2011, Halstead et al. 2017) have demonstrated that ecologically relevant concentrations of atrazine are capable of increasing snail populations, the source of trematode infections to both amphibians and humans. Given the controversy surrounding the effects of atrazine on amphibians, I now follow this reanalysis with a timeline of some of the most salient events in the history of the atrazine-amphibian controversy.

**Fig. 1.**
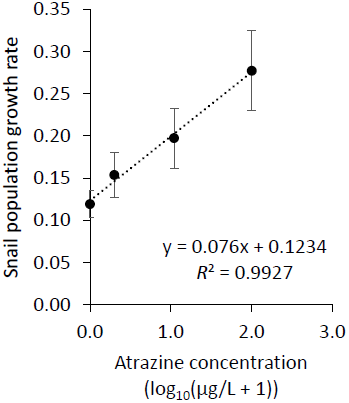
The population growth rate of snails in outdoor mesocosms containing 0, 1, 10, 100 μg/L of atrazine in the Syngenta-funded study by Baxter et al. (2011). Growth rates are calculated for the exponential phase until a carrying capacity or decline in growth occurred. The 30 μg/L treatment was excluded because it did not show the same spatial blocking patterns in dissolved oxygen as the other treatments (see Rohr et al. 2012).

## A Timeline of the Atrazine-Amphibian Controversy

### The early years

The atrazine-amphibian controversy all began in 1998, when Dr. Tyrone Hayes, a biology professor at the University of California at Berkeley, was hired by EcoRisk Inc., the consulting company that hired several academic scientists to study atrazine on behalf of Syngenta Crop Protection, Inc. The contract covering Dr. Hayes' research, and that of many of the other scientists Syngenta and EcoRisk hired, made clear that Syngenta retained final say over what and whether the scientists could publish. In November of 2000, Hayes quit Syngenta because the company supposedly prevented him from publishing his research showing that levels of atrazine, below the drinking water standard of 3 ppb set by the EPA, caused hermaphroditism and reduced the larynx size of male frogs. According to Hayes, Syngenta and Ecorisk tried to keep him working on atrazine privately, offering him as much as $2-million in lab support under the auspices of a start-up company owned by his wife. Hayes refused the offer and began replicating the Syngenta-funded studies using independent funds. Soon after breaking ties with EcoRisk and Syngenta, Hayes claims that Syngenta threaten to pull all of UC Berkeley’s pharmaceutical and medical funding provided by Syngenta’s sister company Novartis Inc. if they tenured Hayes. Despite the ostensible threat, UC Berkeley did eventually tenure Hayes.

At a similar time, in the early 2000s, Krista McCoy began a PhD program at the University of Florida in the laboratory of Dr. Tim Gross, a paid EcoRisk consultant. Krista began a mesocosm study examining the effects of atrazine on amphibians. She came in one morning to discover that Gross had ordered the University of Florida’s physical facilities to pick up McCoy’s mesocosms with a forklift and move them all directly under the roof line of a large nearby building. McCoy was convinced that the mesocosms were moved so they would receive the entire roof’s worth of rain, unrealistically diluting the atrazine. McCoy suspended her atrazine work and switched to laboratory of Dr. Louis Guillette, who confirmed McCoy’s story. Dr. Gross was eventually let go from the University of Florida.

In February of 2002, Dr. Jason Rohr, was hired at the University of Kentucky to study the effects of atrazine on amphibians. In April of 2002, soon after Rohr was hired, Dr. Hayes published his studies repeating the work he originally conducted for Syngenta (Hayes et al. 2002b). This work was published in the prestigious *Proceedings of the National Academy of Sciences of the United States of the America* (*PNAS*), and showed that very low levels of atrazine reduced the larynx size of male frogs and caused male frogs to develop female gonads. According to Hayes, editors at the prestigious journal *Nature* then commissioned him to write a follow-up article on field patterns of atrazine and amphibian gonadal abnormalities that was published in *Nature* in October of 2002 (Hayes et al. 2002a).

In November of 2002, attorneys associated with the Center for Regulatory Effectiveness, the Kansas Corn Growers Association, and the Triazine Network (which receive financial support from Syngenta), argued that Hayes’ studies did not conform with the 2001 Data Quality Act, which prohibits federal agencies from using scientific findings for which there are no established standards. Their petition successfully blocked the US Environmental Protection Agency (EPA) from considering Hayes' work and atrazine was re-registered for use in October of 2003. Ironically, this was the same month that the European Union banned atrazine because of ubiquitous and unpreventable water contamination (Sass and Colangelo 2006). Because of this petition and the Data Quality Act, the EPA had to revise its Environmental Risk Assessment policies, so that hormone disruption would not be a legitimate reason for restricting the use of a chemical until “appropriate testing protocols have been established.” (Sass and Devine 2004). The Data Quality Act has been used widely by industry to block unwanted regulations and as a broader assault on academic freedom (Michaels and Monforton 2005, Rohr and McCoy 2010a).

Since leaving EcoRisk and Syngenta in 2000, the relationship between Hayes and Syngenta representatives became further strained. In 2003, Hayes received a job offer from Duke University and made a second visit to the campus. Duke University is close to Syngenta Crop Protection headquarters in Greensboro, North Carolina and to Syngenta’s research facility in Research Triangle Park, North Carolina. Once Syngenta found out about the offer, they contacted administrators at Duke. Soon after, Duke University withdrew the offer to Hayes. According to subpoenaed documents revealed in a lawsuit (see below), by interfering with Hayes’ job offer, Syngenta was attempting to protect their reputation in their local community and among their employees (Howard 2013a). In October of 2003, The Chronicle of Higher Education published a lengthy article on the damaged relationship between Hayes and Syngenta and the price Hayes had to pay to publish his research (Blumenstyk 2003).

### Tensions rise

In November of 2003, there was an organized oral session on the effects of atrazine on amphibians at the North American Society for Environmental Toxicology and Chemistry meetings in Austin, TX. In attendance were Hayes, Rohr, several EPA representatives, and Syngenta-and EcoRisk-funded scientists, including Ronald Kendall, the head of the EcoRisk panel coordinating the investigation of atrazine for Syngenta, and Keith Solomon, a long-time Syngenta-funded academic. There was standing room only. Much to the surprise of all, Hayes presented no data. Rather, he presented only emails ostensibly incriminating the EPA and Syngenta associates of colluding to ensure the re-registration of atrazine. Rohr presented his first talk ever on atrazine immediately after Hayes, quite surprised and intimidated by what just transpired. Because of Hayes’ bold presentation, SETAC had to hire extra security for their North American Meetings for several years to come. In December of 2004, Hayes continued to keep a target on Syngenta, publishing an article with colleagues in *BioScience* reporting that the single best predictor of whether or not the herbicide atrazine had a significant effect in a study was whether Syngenta funded it (Hayes 2004). That result was highly significant by the usual statistical measures. In 2005, in a lawsuit against the EPA, the Natural Resources Defense Council obtained documents revealing that agency officials met privately with Syngenta more than 40 times while evaluating the toxicity of atrazine (Slater 2012).

Hayes and colleagues’ assault on Syngenta was getting intense and Syngenta began to even more vigorously fight back. In 2005, Syngenta began spending millions on a Hayes ‘smear campaign’ where they came up with a long list of methods for discrediting him, such as “have his work audited by 3rd party,” “ask journals to retract his science,” “set trap to entice him to sue,” “investigate funding,” and “investigate wife” (Howard 2013a, Aviv 2014). They even bought the worldwide web search results for his name so they could better control what the public read about Hayes and atrazine (Howard 2013a, Aviv 2014). Although Hayes suspected much of this, it was not verified until this smear campaign became public in 2012 when thousands of Syngenta documents were subpoenaed in a lawsuit (Howard 2013a) (see below).

Rohr became a bigger target than before in 2008 when he and colleagues published a paper in *Nature* showing that atrazine increased infectious disease risk in a declining amphibian species by reducing frog immunity and increasing its exposure to the pathogen (Rohr et al. 2008c). In November of 2008, in response to an accumulation of papers on atrazine and amphibians, Keith Solomon and colleagues, with financial support from Syngenta, published a paper in Critical Reviews in Toxicology entitled “Effects of atrazine on fish, amphibians, and aquatic reptiles: a critical review” (Solomon et al. 2008). This paper purported to accurately review the effects of atrazine on the behavior, growth, survival, physiology, endocrinology, gonadal morphology, immunity, and infectious disease risk of amphibians. Rohr, a second year professor at the University of South Florida, eagerly read the review paper but did not recall the primary literature the same way as it was described by Solomon et al. (2008). Around the same time, Krista McCoy, an eventual postdoctoral research associate in Rohr’s laboratory, expressed to Rohr that she too did not agree with Solomon et al.’s depiction of the primary literature on atrazine. Hence, Rohr and McCoy collaborated to quantify the inaccurate representations of primary literature in the Syngenta-funded Solomon et al. (2008) article, as well as conduct their own objective meta-analysis of the literature to set the record straight.

While Rohr and McCoy worked on their analyses, the heat on atrazine continued to build. In August of 2009, *The New York Times* investigation found that 33 million Americans were exposed to atrazine through drinking water and, later, data from EPA showed that contamination exceeded the federal limit in 9 out of 10 Midwestern states monitoring it. Several of these water districts reported between 9 and 18 times the federal limit, levels linked to birth defects, premature birth, and low birth weight (Slater 2012).

### The controversy really escalates in 2010

In January of 2010, Hayes et al. published an elegant experiment in *PNAS* (Hayes et al. 2010) where they exposed a laboratory population of all genetically male frogs to low levels of atrazine and showed that these males were both demasculinized (chemically castrated) and completely feminized as adults. Atrazine-exposed genetic males suffered from depressed testosterone, decreased breeding gland size, feminized laryngeal development, suppressed mating behavior, reduced spermatogenesis, and decreased fertility. Additionally, 10% of these males developed into functional females that copulated with unexposed males and produced viable eggs.

During the early months of 2010, Rohr and McCoy completed their assessment of the Solomon et al. (2008) article and set the record straight with their own meta-analysis (Rohr and McCoy 2010b). Rohr and McCoy revealed that the Syngenta-funded review by Solomon et al. (2008) had arguably misrepresented over 50 studies and had 122 inaccurate and 22 misleading statements. Of these 144 seemingly inaccurate or misleading statements, 96.5% appeared to be beneficial for Syngenta in that they supported the safety of the chemical (Rohr and McCoy 2010a). In addition, Solomon et al. (2008) cast doubts on the validity of 94% of the 63 presented cases where atrazine had adverse effects, whereas they almost never criticized the 70 cases where there were no effects of atrazine at environmentally relevant concentrations (Rohr and McCoy 2010a). Rohr and McCoy then conducted a qualitative meta-analysis on the same data analyzed by Solomon et al. (2008) and the general conclusions were the same regardless of whether they excluded studies based on clear quality criteria or included all studies (Rohr and McCoy 2010b). They showed that atrazine regularly disrupted the timing of amphibian metamorphosis, reduced size at or near metamorphosis, altered amphibian motor activity and antipredator behaviors, reduced olfactory abilities, diminished immune function, increased infection end points, and altered aspects of gonadal morphology and function and sex hormone concentrations, but did not directly affect amphibian survival (Rohr and McCoy 2010b). These two studies were submitted as companion papers to *Environmental Health Perspectives*. The meta-analysis was published there (Rohr and McCoy 2010b) but the editor refused to even review the paper on the conflicts of interest, inaccuracies, and bias of the Solomon et al. (2008) paper. After four additional case where editors did not send the paper out for review in many cases fearing the controversy, Rohr and McCoy put a conservation angle on the conflicts of interest paper and published it in *Conservation Letters* (Rohr and McCoy 2010a). There was surprisingly little push back from Syngenta on these papers. In fact, according to subpoenaed documents, Syngenta representatives prepared their funded scientists on how to respond to difficult questions about these studies, describing the meta-analysis as a “rigorous and comprehensive review”.

In July of 2010, Danielle Ivory of the Huffington Post Investigative Fund reported that fewer than 20% of the papers the EPA relied upon in its past decision-making on atrazine were peer-reviewed. Additionally, at least half were conducted by scientists with a financial stake in atrazine (Ivory 2009, 2010). This investigation raised additional concerns over the decision-making process on the safety of atrazine.

In August of 2010, Syngenta struck back against Hayes. They released a 102 page PDF file documenting offensive and potentially embarrassing emails sent by Hayes to Syngenta representatives over the years. In these emails, Hayes had used profanities and sexual taunts, and aggressive, salacious, lewd, and insulting language. The NY Times wrote a story about the emails (Schor 2010) and republished the PDF file (http://www.atrazine.com/amphibians/combined_large_pdf-r-opt.pdf). The emails were also covered by *Nature* (Dalton 2010). These emails made it clear that the unprofessionalism and questionable decision making was occurring by both parties. Dashka Slater provided the following quote in her Mother Jones article to describe these emails “His [Hayes] irreverence had always been an asset, attracting attention to atrazine just as Rachel Carson's impassioned lyricism drew attention to DDT. But now irreverence had tipped toward irrationality.” (Slater 2012). Based on these emails, Syngenta issued a formal ethics complaint filed at the University of California Berkeley. The university's chief counsel ruled that no ethics violation had occurred but admonished both sides to behave.

In 2010, a class-action lawsuit against Syngenta picked up steam. The lawsuit, originally filed in 2004 by Holiday Shores Sanitary District, grew and eventually included more than 1,000 community water systems in Illinois, Missouri, Kansas, Indiana, Iowa and Ohio. The lawsuit, led by Stephen Tillery of the law firm Korein Tillery, LLC, was brought because Midwestern water treatment facilities often could not get atrazine concentrations in their drinking water below the US EPA maximum contaminant level deemed safe for human consumption (3 ppb). Rohr passed on testifying in the case (see below), whereas Hayes did testify and Stephen Tillery stated that Hayes’ work gave them the scientific basis for the lawsuit.

In 2012, the class-action lawsuit filed in 2004 by Holiday Shores Sanitary District that grew into a class action lawsuit with more than 1,000 community water systems in Illinois, Missouri, Kansas, Indiana, Iowa and Ohio vigorously continued until the integrity of an important witness for Korein Tillery (someone other than Hayes) was questioned after illicit behaviors were allegedly uncovered. Soon after, Tillery and associates settled the suit but Syngenta denied all wrongdoing and did not claim any liability. Syngenta paid $105 million to reimburse more than a thousand Midwestern water utilities for the cost of filtering atrazine from drinking water. When lawyer fees were removed, this amounted to well under $100,000 per water treatment plant.

Stephen Tillery and another lawyer from his office flew from Illinois to Rohr’s office to recruit him as an expert scientist for the case. They offered to pay Rohr generously for his services. Before jumping at the opportunity, Rohr queried Tillery. Rohr asked whether the atrazine problem was not at least partially an issue of how much atrazine was applied in the Midwestern US and that most of the water treatment plants there lacked modern carbon filtration systems necessary to remove the atrazine. Tillery confirmed that this was a major source of the problem. Rohr then made it clear that Syngenta could not control who buys their product, where they apply it, how much they apply, or whether water treatment plants have adequate carbon filtration systems. Hence, Rohr questioned whether the atrazine problem in the Midwestern US was an EPA enforcement issue and whether the EPA, not Syngenta, should be sued. Tillery agreed with all the logic but claimed that he could not sue the EPA. Rohr, however, made it clear to Tillery that he could sue the EPA because the Natural Resource Defense Council sues the EPA all the time; law firms just cannot sue the EPA for money. Rohr politely declined the offer to be an expert witness, worried that the lawsuit was misdirected at the entity with the deepest pockets.

In November of 2010, Rohr gave a seminar on atrazine at Illinois State University the year after Hayes gave his seminar there. Illinois State University is not far from Syngenta’s US headquarters in Illinois. In preparation for the seminar, Illinois State University’s Department of Biology hired a security guard to staff the talk. Syngenta sent an attorney in a three piece suit that was frisked by the security guard to check for recording devices. The attorney took copious notes throughout the talk.

After the flurry of papers published by Rohr’s laboratory between 2008 and the beginning of 2011 on atrazine and chlorothalonil, two pesticides produced by Syngenta, Rohr started receiving pushback from the Director of the facility where Rohr conducted his outdoor tank (mesocosm) experiments on agrochemicals. Rohr collaborated with Dr. Steven Johnson of the University of Florida’s Gulf Coast Research and Education Center (GCREC), which was just 45 minutes from the University of South Florida where Rohr is employed. This collaboration allowed Rohr to have his tank facility at GCREC. The Director of the GCREC’s wife worked for Syngenta at the time. Rohr and Johnson did not face any challenges until they started publishing their findings. Johnson surprisingly found out that he was being forced out of the GCREC back to the main campus of the University of Florida in Gainesville. Given that Johnson was no longer at the GCREC, the Director of the GCREC told Rohr he also had to leave because his collaborator was no longer at the facility. Rohr resourcefully looked to find another collaborator at the GCREC so he could continue his work there. He found two faculty members other than Johnson that wanted to collaborate but the Director of the GCREC blocked both collaborations and eventually told all GCREC faculty that they could not collaborate with Rohr or conduct any pesticide-related toxicological research, stifling their academic freedoms. It took Rohr several years to find a location and setup another tank facility, a major impediment to his ecotoxicological research on atrazine and other agrochemicals.

### Revelations from the lawsuit and the EPA scientific advisory panel

The most important outcome of the lawsuit was not the settlement but the Syngenta documents that became “unsealed” by the Madison County Circuit Court in response to a Freedom of Information Act request by the courageous investigative reporting conducted by Clare Howard of 100Reporters. The 1,000 or so pages of memos, notes, and e-mails that Clare received exposed Syngenta’s tactics and efforts to conceal and discredit the science on atrazine. They revealed that Hayes was not paranoid after all and that Syngenta was indeed behind a campaign to smear him and his reputation. The subpoenaed documents revealed that one of the company’s strategies had been to “*purchase ‘Tyrone Hayes’ as a search word on the internet, so that any time someone searches for Tyrone’s material, the first thing they see is our material*.” Syngenta later also purchased the phrases “amphibian hayes,” “atrazine frogs,” and “frog feminization” and searching online for “Tyrone Hayes” for years until the settlement brought up an advertisement that said, “Tyrone Hayes Not Credible.” The documents revealed that Syngenta invested in a multi-million dollar campaign to protect atrazine profits, which included hiring a detective agency to investigate scientists on a federal advisory panel, looking into the personal life of a judge, and discrediting and distracting Hayes. These documents also listed other strategies directed at Hayes, such as “*commissioning a psychological profile*”, “*have his work audited by 3rd party*”, “*ask journals to retract*”, “*set trap to entice him to sue*”. “*investigate funding*”, “*investigate wife*”, “*tracking him at speaking engagements*”, “*baiting him through emails*”, and interfering with Hayes’ job offer at Duke. Syngenta would send representative to talks Hayes gave to question and embarrass him. Rohr too had similar experiences. The documents also revealed that Syngenta kept a list of 130 people and groups it could recruit as experts, including academics, without disclosing ties to the company (see https://www.documentcloud.org/documents/686401-100reporters-syngenta-clare-howard-investigation.html for the list). It often paid members of this group to write opeds and other articles. According to Jayne Thompson from Jayne Thompson & Associates, a public relations firm hired to work on the Syngenta campaign, “*These are great clips for us because they get out some of our messages from someone who comes off sounding like an unbiased expert. Another strength is that the messages do not sound like they came from Syngenta*.” Clare Howard summarizes her investigative reporting on these documents in a ground breaking article published in June of 2013 in a 100Reporters (Howard 2013a). Unfortunately, these documents did not get considerable press until February of 2014 when the more well-known magazine *The New Yorker* released an article on atrazine, Hayes, and the uncovered Syngenta documents (Aviv 2014). The original *New Yorker* article inexplicably did not acknowledge any of Clare’s seminal investigative work (Aviv 2014).

Ironically, as the class-action lawsuit was being settled, so too were the policy decisions on the safety of atrazine to amphibians. The EPA had convened a scientific advisory panel to assess the effects of atrazine on amphibians and originally offered Rohr to be a member on this panel. The EPA then rescinded this offer because Rohr’s work would serve too prominently in panel discussions. The EPA then recruited Dr. Michelle Boone to take Rohr’s place. The USEPA concluded that “*exposure to atrazine at concentrations ranging from 0.01 to 100 [milligrams per liter] had no effect on Xenopus laevis [an amphibian species] development (which included survival, growth, metamorphosis, and sexual development)*” (p. 60) and that the “*level of concern for effects on aquatic plant communities… was lower than the atrazine concentration observed to produce significant direct or indirect effects on invertebrates, fish, and amphibians*” (USEPA 2012), which would eliminate further assessments of atrazine’s impacts on amphibians despite significant effects at these concentrations in other studies. The EPA’s conclusion that atrazine does not adversely affect amphibians, however, was based on a single published study that was funded by Syngenta, despite there being hundreds of studies on atrazine. Several members of the scientific advisory panel argued that a decision on the safety of atrazine should not be made based on any single study, especially one funded by the company with a financial stake in the product. In fact, after finishing her work on the scientific advisory panel, Boone collaborated with Rohr and other colleagues to write articles denouncing the use of a single industry-funded study to evaluate the adverse effects of atrazine, or for that matter, any chemical (Boone et al. 2014, Boone and Rohr 2015).

In April of 2014, Rohr, Hayes, and Solomon appeared on 16×9, a Canadian National Television primetime news show similar to 60 Minutes, 20/20, or Dateline in the US (http://globalnews.ca/video/1252483/full-story-pesticide-peril). The story was produced by Gil Shochat and summarizes the story of the amphibian-atrazine controversy. It also offers an interview with a former Syngenta staffer who describes Syngenta’s internal strategies for discrediting scientists.

### Three additional surprising twists

In August of 2014, UC Berkeley shut down Hayes’ amphibian research because Hayes did not have sufficient funds to pay for his vertebrate animal care. Hayes claimed that his lab fees had gone up by 295% since 2004, while fees for his colleagues at UC Berkeley had risen by only 15% (Howard 2013b). The director of the office of laboratory-animal care apparently provided evidence that Hayes was being charged according to standard campus-wide rates that increased for most researchers in recent years. Nevertheless, Hayes recruited Stephen Tillery to represent him in a lawsuit against UC Berkeley claiming that his vertebrate animal care fees were essentially preventing him from doing his job. The status of this case is currently unclear.

In 2016, after the damaging press of the unsealed court documents, Syngenta announced that it was set to be acquired by Chinese state-owned ChemChina (Spegele and Chu 2016). However, Syngenta and ChemChina missed the European Union’s deadline for submission of antitrust remedies (Blackstone and Drozdiak 2016), raising questions regarding whether they will be able to satisfactorily deal with the antitrust concerns. It remains unclear whether the deal will happen and if the tactics of Syngenta will change if the sale occurs.

In May 2007, the Center for Biological Diversity filed a lawsuit against the EPA for violating the Endangered Species Act by registering and allowing the use of many pesticides without determining whether the chemicals jeopardized endangered species in the San Francisco Bay. A federal court then signed an injunction, imposing interim restrictions on the use of 75 pesticides in the Bay Area. This in turn required that the EPA formally evaluate the effects of those chemicals on endangered species. In June of 2015, the EPA announced that it would analyze the effects of glyphosate (active ingredient in the herbicide Roundup) and atrazine on 1,500 endangered plants and animals (Beyond_Pesticides 2015). Likely as a result of this court order, the EPA re-evaluated atrazine and, apparently in error, released the report online in April of 2016, a presidential election year. This sparked criticism from Syngenta and U.S. lawmakers (Polansek 2016).

In perhaps the most surprising twist of all, the EPA completely reversed its position on the safety of atrazine in this “inadvertently” released reassessment of atrazine. Despite the EPA concluding that atrazine was safe for over four decades, the new and refined risk assessment (Farruggia et al. 2016) states the following: “*Based on the results from hundreds of toxicity studies on the effects of atrazine on plants and animals, over 20 years of surface water monitoring data, and higher tier aquatic exposure models, this risk assessment concludes that aquatic plant communities are impacted in many areas where atrazine use is heaviest, and there is potential chronic risk to fish, amphibians, and aquatic invertebrates in these same locations. In the terrestrial environment, there are risk concerns for mammals, birds, reptiles, plants and plant communities across the country for many of the atrazine uses. EPA levels of concern for chronic risk are exceeded by as much as 22, 198, and 62 times for birds, mammals, and fish, respectively. For aquatic phase amphibians, a weight of evidence analysis concluded there is potential for chronic risks to amphibians based on multiple effects endpoint concentrations compared to measured and predicted surface water concentrations… average atrazine concentrations in water at or above 5 μg/L for several weeks are predicted to lead to reproductive effects in fish, while a 60-day average of 3.4 μg/L has a high probability of impacting aquatic plant community primary productivity, structure and function*.” It remains unclear why the EPA changed its opinion on the safety of atrazine, how the report was “inadvertently released”, and what its consequences will be. However, if there is one thing that I have learned about the science and policy decisions on atrazine, it is to expect the unexpected!

